# Language processing in brains and deep neural networks: computational convergence and its limits

**DOI:** 10.1101/2020.07.03.186288

**Authors:** Charlotte Caucheteux, Jean-Rémi King

## Abstract

Deep Learning has recently led to major advances in natural language processing. Do these models process sentences similarly to humans, and is this similarity driven by specific principles? Using a variety of artificial neural networks, trained on image classification, word embedding, or language modeling, we evaluate whether their architectural and functional properties lead them to generate activations linearly comparable to those of 102 human brains measured with functional magnetic resonance imaging (fMRI) and magnetoencephalography (MEG). We show that image, word and contextualized word embeddings separate the hierarchical levels of language processing in the brain. Critically, we compare 3,600 embeddings in their ability to linearly map onto these brain responses. The results show that (1) the position of the layer in the network and (2) the ability of the network to accurately predict words from context are the main factors responsible for the emergence of brain-like representations in artificial neural networks. Together, these results show how perceptual, lexical and compositional representations precisely unfold within each cortical region and contribute to uncovering the governing principles of language processing in brains and algorithms.

## 1 Introduction

Convergent evolution – when distantly related species (e.g. bats and birds) develop similar structures or functions (i.e. wings) – is often critical to reveal the principles that guide the variety life forms (i.e. controlling aerodynamics with minimal energy). Convergence can be investigated in artificial agents too: “deep” artificial neural networks have recently made substantial progress in harnessing abilities considered uniquely human (4; 5; 6). In language, in particular, deep nets demonstrate unprecedented completion, translation, and summarization abilities (7; 8; 9; 10). Do these algorithms process language similarly to the human brain? Does this similarity directly depend on their training? In sum, is there, in the domain of language processing, a computational convergence between brains and deep neural networks?

These questions are all-the-more challenging that the neurobiology of language remains in its infancy. Previous studies showed that reading depends on a cortical hierarchy originating in the primary visual cortex (V1), propagating within the visual word form area (in the fusiform gyrus, where letter strings are recognized) and reaching the angular gyrus, the anterior temporal lobe and the middle temporal gyrus – associated with lexical understanding (11; 12; 13; 14; 15). This hierarchical pathway and a parallel motor route (13) together connect to the inferior frontal gyrus, where Broca area presumably performs key compositions, like syntax (12; 13; 16; 17; 18). However, the precise nature, format, and dynamics of such lexical and compositional representations is still unclear (18; 19; 13)

This challenge has been partly addressed with Natural Language Processing (NLP) algorithms. For example, word embeddings – high dimensional dense vectors shaped to predict the average lexical neighborhood (20; 21; 22; 23) – have been shown to linearly correlate with the brain responses elicited by words presented either in isolation (24; 25; 26) or within narratives (27; 28; 29; 30; 31; 32). More recently, *contextualized* word embeddings improved such correlations, especially in the prefrontal, temporal and parietal cortices (33; 34; 35).

However, these studies focused on a handful of heterogeneous pretrained models, typically varying in dimensionality, architecture, training objective and corpora. Yet, random embeddings can capture relevant dimensions (1; 2), and consequently lead a network to significantly correlate with brain activity. Consequently, it is unclear whether deep neural networks trained on language modeling systematically (1) converge to, (2) anecdotally correlate with, or (3) even diverge away from brain representations during their training (Figure 1 A).

**Figure 1:**
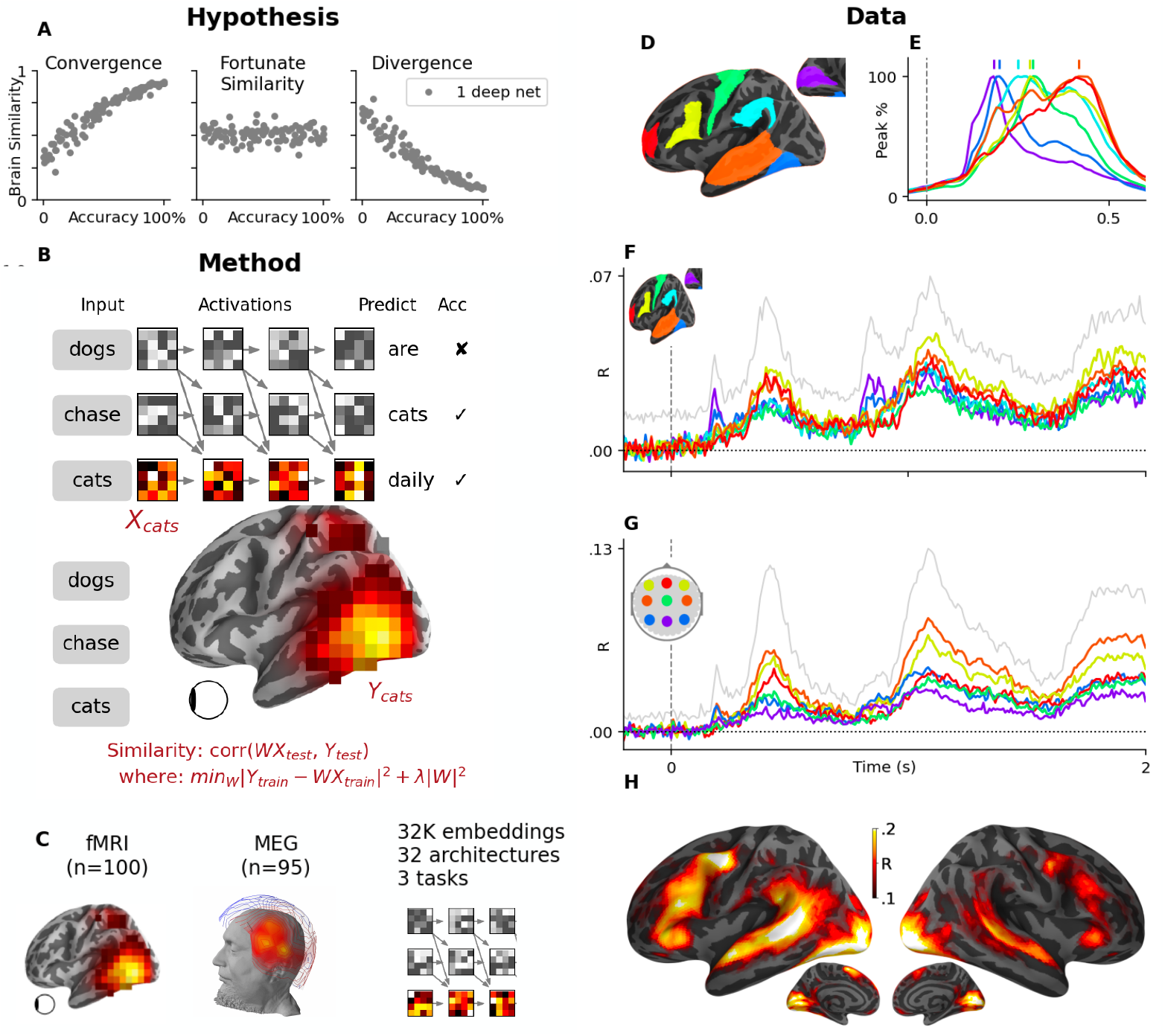
Testing the convergence hypothesis between artificial neural networks and the human brain. **A**. Artificial neural networks would be considered to *converge* to brain-like computations if and only if training consistently increases the similarity between their activations and those of the brain, when input with the same stimuli. Such similarity may be observed, because high dimensional embeddings can contain relevant components by chance (1; 2). Each dot, in the panels, represents one hypothetical embedding: i.e. the activations of a single neural network trained with a fixed amount of data. **B**. The network depicts a 4-layer causal transformer trained to predict words from a preceding context. The similarity between such transformer and the brain is assessed with a linear regression *W* (1) predicting brain responses *Y* from the model’s activations(*X* in response to the same stimuli and (2) evaluated with a correlation between the predictions and true brain responses to held-out sentences *Y*_*test*_ (3). **C**. Using 100 fMRI recordings and 95 MEG recordings, we compare 32,400 embeddings, derived from 32 architectures trained on 3 distinct tasks and evaluated on 100 training steps. **D**. Grand average MEG source estimates to word onset (t=0) for 7 regions typically associated with reading (V1: purple, M1: green, fusiform gyrus: dark blue, supramarginal gyrus: light blue, superior temporal gyrus: orange, infero-frontal gyrus: yellow and fronto-polar gyrus: red), normalized to their peak response. Vertical bars indicate the peak time of each region. The full (not normalized) data is displayed in Video 1. **E**. MEG noise ceilings, approximated by predicting brain responses of a given subject from those of all other subjects. Colored lines depict the mean noise ceiling in each region of interest. The grey line depict the best noise ceiling across sources. **F**. Same as (D) in sensor space. **G**. Noise ceiling estimates of fMRI recordings.

Here, we evaluate whether the activations of 3,600 neural network embeddings linearly correlate with functional magnetic resonance imaging (fMRI) and source-localized magneto-encephalography (MEG) of 102 subjects reading 400 distinct and isolated sentences (36). We then evaluate how each functional and architectural properties of the networks predict their similarity with brain responses.

Our study provides three main contributions. First, we confirm that pretrained neural networks, and their middle layers in particular, linearly correlate with a variety of brain responses to words, even when those are presented within isolated sentences. Second, we show how these networks help track and isolate the sequential generation of perceptual, lexical and compositional representations within each of these cortical regions. Finally, we show that the convergence of deep language models towards brain-like responses is (1) limited to specific layers and (2) predominantly driven by their ability to accurately predict words from context.

## 2 Results

### 2.1 Average and single-trial fMRI and MEG responses to reading

We first aim to identify where and when cortical neurons are activated during a reading task. As expected (18; 12; 37; 13), the rapid serial visual presentation of words elicited responses in a distributed bilateral network, including the primary visual cortex, the left fusiform gyrus, the supra marginal and superior temporal cortices, as well as the motor, premotor and infero-frontal areas (Figure 1). MEG source reconstruction further clarifies the dynamics of this network: on average, word onset elicited a sequence of brain responses originating in V1 around *≈*100 ms and continuing within the left posterior fusiform gyrus around 200 ms, the superior and middle temporal gyri, as well as the pre-motor and infero-frontal cortices between 150 and 500 ms after word onset (Figure 1, Video 1).

What proportion of these brain responses can be accounted for by the specific content and form of each word in each sentence? To address this issue, we trained, for each subject separately, a “noise-ceiling” model across subjects. Specifically, for each recording of each subject and each sentence *Y*_*train*_, we trained a linear model *W* from the recordings of all other subjects who read the same sentence *X*_*train*_. Using a cross-validation scheme across sentences, we then evaluated the Pearson correlation *R* between (1) the true brain responses of subject *Y*_*test*_ and (2) the predicted brain responses *Ŷ*_*test*_ = *W X*_*test*_. This procedure could be thought of as approximating an optimal black box: a one-hot encoder of brain responses is fitted to each element of a unique sentence. The results are summarized in Figure 1 F-H. As expected, noise ceiling estimates peaked within the well-known language network (38) were substantially lower in MEG (especially in source space) than in fMRI. For example, fMRI noise ceilings reached, on average, *R* = 0.129 (*±*0.004 SEM across subjects) in the superior temporal gyrus whereas MEG noise ceilings peaked at *R* = 0.069 *±* 0.001.

### 2.2 Image, word and compositional embeddings correlate with different parts of brain activity

Following previous studies (39; 40; 41; 3; 26; 27; 28; 29; 31; 33; 34; 35; 18), we evaluated whether the activations of (1) a visual neural network, (2) a word embedding and (3) a contextualized word embbeding can linearly predict MEG and fMRI responses to words presented in isolated sentences.

For the image embedding, we input an image of each word to a deep convolutional neural network (CNN) trained on character recognition (42) and extracted the activations of the last layer. Similarly, we input the sentences that the subjects read to a 13-layer transformer trained on language modeling and extracted the 128-dimensional activations of the first and middle layers to retrieve word and compositional embeddings, respectively.

For each embedding *X*, we trained and evaluated with cross-validation, the ability of a linear mapping *W* to predict brain responses *Y*. Figure 2 and Video 2 summarize the corresponding MEG and fMRI correlation scores. For clarity, Figure 2C and the video plot the gain in MEG scores obtained by word embeddings (compared to visual embeddings) and by compositional embeddings (compared to word embeddings).

**Figure 2:**
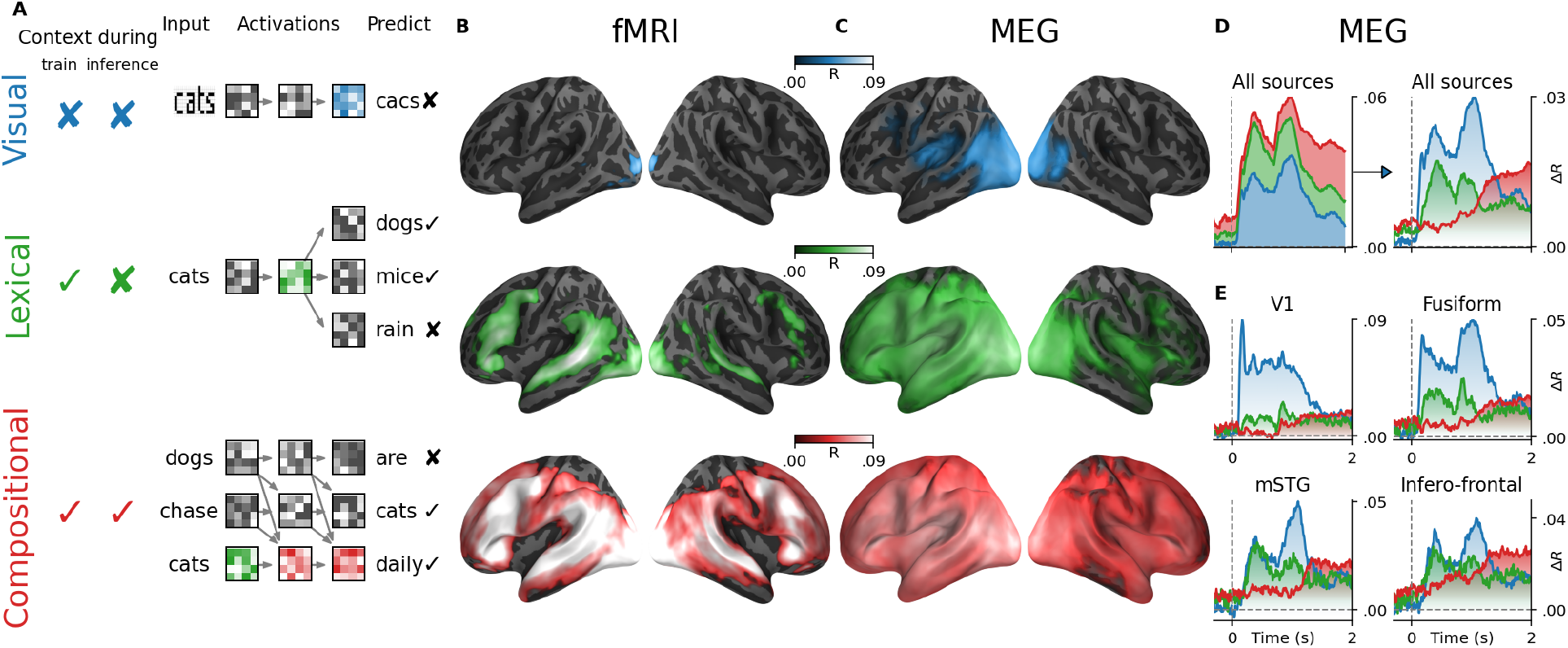
Encoding of hierarchical representations. **A**. Visual, lexical, and compositional representations can be isolated from a convolutional neural networks trained on character recognition (top, blue), a word embedding (middle, green) and the middle layer of a transformer trained on language modeling (bottom red), because each of these embeddings accesses word context variably during training and/or inference. **B**. Mean (across subjects) fMRI encoding scores obtained with the convolutional neural network (top, blue), the word embedding (middle, green) and the transformer (bottom, red). **C**. Mean MEG encoding scores averaged across all time samples. **D**. Mean MEG encoding scores averaged across all sensors (left) and the corresponding gains (i.e. green: [word embedding] - [visual embedding]; red: [compositional embedding] - [word embedding]). **E**. Mean gains in MEG encoding scores averaged within four regions of interest. For a whole-brain depiction of the gains in encoding scores, see Video 2. Shaded regions highlight significant scores across subjects (*p <* 10^*−*2^after FDR correction).

The brain scores of image embedding peaked in the early visual cortex (V1), both for fMRI (*R* = 0.022 ± 0.003, *p <* 10^*−*11^ across subjects) and MEG source estimates (*R* = 0.008 ± 0.002, *p <* 10^*−*3^, for t=100ms). These MEG scores peaked around 100 ms in V1, and rapidly propagated to higher-level areas. Word embeddings peaked around 400 ms and were primarily distributed over the left temporal (*R* = 0.052 ± 0.004, *p <* 10^*−*15^) and frontal cortices (*R* = 0.053 ± 0.003, *p <* 10^*−*15^). Finally, compositional embeddings mainly improved brain scores in the superior temporal gyrus (Δ_*R*_ = 0.012 ± 0.001, *p <* 10^*−*16^), the angular gyrus (Δ_*R*_ = 0.010 ± 0.001, *p <* 10^*−*16^), the infero-frontal cortex (Δ_*R*_ = 0.016 ± 0.001, *p <* 10^*−*16^) and the dorsolateral prefrontal cortex (Δ_*R*_ = 0.012 ± 0.001, *p <* 10^*−*13^). These effects were mostly lateralized (between left and right, Δ_*R*_ = 0.010 ± 0.001, *p <* 10^*−*14^). The gain in MEG-scores obtained with compositional embeddings appears to be mainly driven by brain responses after *≈* 1sec, and is observed in a large number of bilateral brain regions (Figure **??**C-D).

Around this time window, brain areas outside the language network, such as area V1, appeared to be better predicted by word and compositional embeddings than by visual ones (e.g between visual and word in V1: Δ_*R*_ = 0.016 ± 0.002, *p <* 10^*−*10^). These effects could thus reflect feedback activity (43)and explain why the corresponding fMRI responses are better accounted for by word and compositional embeddings than by visual ones.

Overall, these results confirm that the activations of three typical deep neural networks trained on image, word or sentence processing linearly mapped onto brain responses to the same input. In addition, these results allow us to track, with an unprecedented spatio-temporal precision, the hierarchical generation of visual, lexical and compositional representations in each cortical region (Video 2).

### 2.3 The middle layers of trained language models best predict brain responses independently of their tasks and architectures

To what extent are the above correlations representative of the similarity between brains and deep neural networks used in natural language processing ? To address this issue, we analysed a variety of Transformers – state-of-the-art feedforward networks that rely on an attention mechanism to combine words into meaning (7; 8). Specifically, we implemented 32 architectures (from 4 to 12 layers, each varying from 128 to 512 dimensions, and each benefiting from 4 to 8 attention heads), trained each of them on two distinct tasks (“causal” language modeling or “masked” language modeling), assessed the extent to which their activations linearly predicted fMRI and MEG responses, and evaluated how their architectural and functional properties impacted the brain scores.

Brain scores mainly varied as a function of the relative position of each extracted layer (Figure. 3). Specifically, middle layers systematically outperformed output (fMRI: Δ_*R*_ = 0.011 ± 0.001, *p <* 10^*−*18^, MEG: Δ_*R*_ = 0.003 ± 0.000, *p <* 10^*−*13^) and input layers (fMRI: Δ_*R*_ = 0.031 ± 0.001, *p <* 10^*−*18^, *MEG* : Δ_*R*_ = 0.009 ± 0.001, *p <* 10^*−*17^). This effect was consistent across 32 architectures and two training tasks, cf. Figure 3.

**Figure 3:**
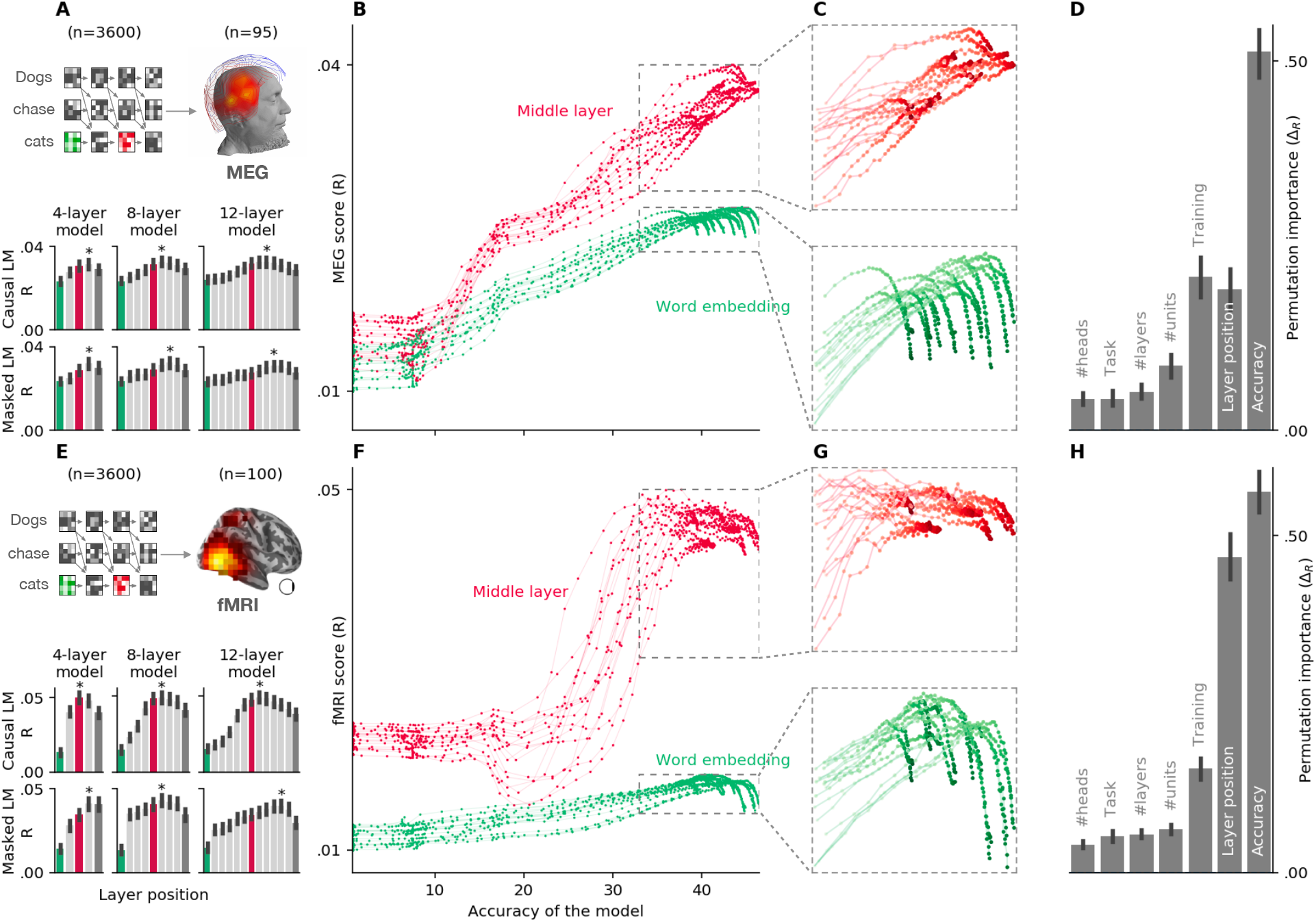
Only middle layers of language models consistently converge to brain representations. **A**. Bar plots display the MEG encoding score (averaged across time and channels) of 6 representative transformers varying in tasks (CLM vs MLM) and depth (4-12 layers). The green and red bars correspond to the word-embedding and the best layers respectively. **B**. MEG scores (mean across subjects, time and channels) of each of the 16,200 embeddings extracted from 1,800 causal deep networks (dots), separately for the input layer (word embedding, *l* = 0, green) and the middle layer (red). On the x-axis, the network’s language modeling performance (top-1 accuracy to predict the next word). Each line corresponds to one architecture. Each dot correspond to their embedding extracted at a specific training step. **D**. Same as (C), zooming on the best-performing neural networks (accuracy >35). **E**. Results of the feature importance analysis. Each bar indicates how much each model variable contributes to making the activations of the artificial neural network similar to the MEG responses (cf. Methods). **F-J**. Same as above, but the brain-score is evaluated on the fMRI recordings of subjects (as opposed to the MEG recordings). Error bars are the 95% confidence intervals of the scores’ distribution across subjects.

To assess how each model property explained brain scores, we implemented a permutation importance analysis across models with a random forest ((44)), for each subject independently (cf. Methods for more details). Overall, the relative position of the layers explained 81.5 ± 1.2% of fMRI and 70.2 ± 1.4% of MEG scores, whereas architecture and task variables accounted for less than 17% of them. In sum, this analysis confirms that the results described in section 2.2 are representative of language models.

### 2.4 Only middle layers converge towards brain responses

The above similarities between brain responses and artificial neural networks result from the analysis of trained networks. Yet, random neural networks can contain relevant features (1; 2), and can thus significantly predict brain responses. To test whether training leads neural networks to (1) converge to, (2) fortunately correlate with or even (3) diverge away from brain-like solution, we applied brain score analyses for each artificial neural network frozen at 100 different training steps. We then tested whether the similarity between their activations and brain responses consistently increased with their training and language performance (top-1 accuracy at predicting the masked/next word on a test set).

On average, the input layer activations (word embeddings) increased within the 100K first training iterations (*≈* 200*M* processed words, one iteration corresponds to one gradient update). However, training ultimately led to a steady decrease, even though the networks continued to improve on their training task (Figure 3): the Pearson correlation between training steps and brain scores after 100K iterations was negative both in MEG (*R* = *−*0.166 ± 0.019, *p <* 10^*−*4^) and in fMRI (*R* = *−*0.106 ± 0.021, *p <* 10^*−*4^).

By contrast, the brain scores of the middle layer (Figure 3) increased with the language modeling accuracy in MEG (*R* = 0.87 ± 0.01, *p <* 10^*−*16^), Figure 3) and fMRI (*R* = 0.82 ± 0.02, *p <* 10^*−*17^). Note that the brain scores of the middle layers stabilized in fMRI slightly before MEG.

Language modeling accuracy varies with other model properties, such as architecture and training parameters. To disentangle how each property contributes to making the model activation more-or-less similar to the brain, we implemented a feature importance analysis for each subject independently (Figure 3 D, H). The results confirmed that language modeling accuracy was the most important factor that leads the network to brain-like solutions: fMRI: Δ*R* = 0.56 ± 0.02, MEG: Δ*R* = 0.51 ± 0.02. This variable was followed in its brain-like contribution by the amount of training the network underwent (fMRI: Δ*R* = 0.15 ± 0.01, MEG: Δ*R* = 0.21 ± 0.01) and the relative layer position of the extracted representation (fMRI: Δ*R* = 0.47 ± 0.02, MEG: Δ*R* = 0.19 ± 0.01, Figure 3 D and H). By contrast, the training task, the dimensionality of the layers, the number of heads, and the total number of layers modestly influenced brain scores (Δ*R <* .08).

Overall, these results suggest that, beyond the marginal effects of the models’ architecture, the middle - but not the input and output - layers of deep language models converge towards brain-like representations.

## 3 Discussion

Do modern algorithms learn to process information in a way that leads them to mimic the computations of the human brain? Following recent achievements (45; 46; 47; 39; 40; 41; 14; 35; 48; 3; 49), we address this challenge on the restricted issue of sentence processing, by evaluating whether the activations of a large variety of neural networks linearly map onto those of 102 human brains, each recorded with MEG and fMRI during an isolated sentence reading task.

We found that the similarity between brains and artificial neural networks mainly depends on the language modeling accuracy of the latter, and is predominantly driven by their middle layers. This result extends recent fMRI (33; 34; 35) MEG (32; 50) and ECoG findings (51; 52), showing that pretrained language models linearly map onto the brain responses to English narratives. Our analyses provide a precise description of the spatio-temporal dynamics underlying linguistic processes. First, we confirm the sequential generation of visual and lexical representations in the fusiform gyrus (Video 2) predicted by reading theories (12; 53; 54). Second, we confirm with isolated and written sentences (and thus devoid of narrative or prosody contours) that word embeddings correlate with a large fronto-temporo-parietal network, and reveal their remarkably sustained effects (24; 33; 35). Third, the compositional representations of deep language models peaked precisely in the brain regions traditionally associated with high-level sentence processing (55; 13; 56). Finally, we subsume previous results based on average responses showing that language composition significantly recruits both hemispheres even though these effects are left-lateralized (57; 58).

Most of the correlation scores reported in the present study are very low. This phenomenon likely results from our use of single sample encoding analysis. These effects, however, appear to be within the range obtained with noise ceiling estimates. While the unusually large number of words and subjects in the analysis allows for a high level of statistical significance, our results emphasize the major limitations of neuroimaging imposed by signal-to-noise ratio.

The convergence observed between brains and deep language models follows a nontrivial pattern. The convergence of middle layers to brain-like representations is partly expected: middle layers have been shown to linearly encode syntactic trees (59) and co-references (60; 61). However, input and output layers ultimately diverged away from brain responses. This result surprised us, especially because word embeddings have been repeatedly used to model brain activity (24; 26; 14). This phenomenon begs the question whether language models learn to *combine* words - as opposed to *represent* them - similarly to humans.

In any case, the convergence of deep language models to brain-like computations is undoubtedly partial. First, modern language models are still far from human-level performance on a variety of tasks such as dialogue, summarization, and systematic generalization (62; 63). Second, the size of their training corpora can be incommensurately larger than what a human may be to read in his or her lifetime (10). Third, the architecture of the popular transformer network (7) is in many ways *not* biologically plausible: while the brain has been repeatedly associated with a *recurrent* predictive coding architecture, where prediction errors are computed at each level of an interconnected hierarchy of recurrent networks (64), transformers are *feedforward* neural networks that access an unreasonably large buffer of words, and only minimize prediction errors at their final “predicted word” layer. In light of these major differences, it is all-the-more remarkable to see that brains and artificial neural networks find a partially common solution to language processing.

Representations were here modeled for each contextualized word independently. However, the precise nature of these representations, both in the brain and in artificial neural networks, is here only coarsely categorized into three hierarchical levels. How the mind builds and organizes its lexicon, how it parses and manipulates sentences, how it plans and memorizes narratives, and perhaps above all, how it learns to achieve all these skills remain open questions. Nevertheless, the present study shows how the comparative study of brains and artificial neural networks may help test the hypothesis, according to which a simple learning objective – predicting words from their context – suffices to explain how the human brain came to form the peculiar structures of language.

## 4 Methods

We assessed the similarity between (i) the activations of deep neural networks and (ii) those of the brain of 100 subjects, measured with magneto-encephalography (MEG) and functional magnetic resonance imaging (fMRI), when the two were input with the same 400 isolated sentences.

### 4.1 Deep Neural Networks

#### 4.1.1 Visual Convolutional Neural Network

To model visual representations, every word presented to the subjects was rendered on a gray 100 x 32 pixel background with a centered black Arial font, and input to a VGG network pretrained to recognize words from images (42), resulting in an 888-dimensional embedding. This embedding was used to replicate and extend previous work on the similarity between visual neural networks activations and brain responses to the same images (e.g. (45; 39; 40)).

#### 4.1.2 Language Transformers

To model word and sentence representations, we trained a variety of transformers (7), input them with the same sentences that the subject read and extracted the corresponding activations from each layer. We always extract activation in a “causal” way: for example, given the sentence ‘THE CAT IS ON THE MAT’, the brain response to ‘ON’ would be solely compared to the activations of the transformer input with ‘THE CAT IS ON’, and extracted from the ‘ON’ contextualized embeddings. Word embeddings and contextualized embeddings were generated for every word, by generating word sequences from the three previous sentences. We did not observe qualitatively different results when using longer contexts. Note that the sentences were isolated, and were not part of a narrative.

In total, we investigated 32 distinct architectures varying dimensionality (*∈* [128, 256, 512]), number of layers (*∈* [4, 8, 12]), attention heads (*∈* [4, 8]) and training task (“causal” language modeling and “masked” language modeling). While “causal” language transformers are trained to predict a word from its previous context, “masked” language transformers predict randomly masked words from a surrounding context. We froze the networks at *≈* 100 training stages (log distributed between 0 and 4,5M gradient updates, which corresponds to *≈* 35 passes over the full corpus), resulting in 3,600 networks in total, and 32,400 word representations (one per layer). Training was early-stopped when the networks’ performance did not improve after 5 epochs on a validation set. Therefore, the number of frozen steps varied between 96 and 103 depending on the training length.

The algorithms were trained using XLM implementation ^1^ (9), on the same Wikipedia corpus of 278,386,651 words extracted using WikiExtractor ^2^ and pre-processed using Moses tokenizer (65), with punctuation. We restricted the vocabulary to the 50,000 most frequent words, concatenated with all words used in the study (50,341 vocabulary words in total). These design choices enforce that the difference in brain scores observed across models cannot be explained by differences in corpora and text preprocessing.

To evaluate the language processing performance of the networks, we computed their performance (top-1 accuracy on word prediction given the context) using a test dataset of 180,883 words from Wikipedia.

### 4.2 Neuroimaging

#### 4.2.1 Protocol

One-hundred and two native Dutch right-handed speakers performed a reading task while being recorded, by Schoffelen and colleagues, with a CTF magneto-encephalography (MEG) and, in a separate session, with a SIEMENS Trio 3T Magnetic Resonance scanner (36).

Words were flashed one at a time with a mean duration of 351 ms (ranging from 300 to 1400 ms), separated with a 300ms blank screen, and grouped into sequences of 9 - 15 words, for a total of approximately 2,700 words per subject. Sequences were separated by a 5s-long blank screen. We restricted our study to meaningful sentences (400 distinct sentences in total, 120 per subject). Twenty percent of the sentences were followed by a yes/no question (e.g. “Did grandma give a cookie to the girl?) to ensure that subjects were paying attention. Sentences were grouped into blocks of five sequences. This grouping was used for cross-validation to avoid information leakage between train and test sets.

#### 4.2.2 Magnetic Resonance Imaging (MRI)

Structural images were acquired with a T1-weighted magnetization-prepared rapid gradient-echo (MP-RAGE) pulse sequence. The full acquisition details, available in (36), are here summarized for simplicity: TR=2,300 ms, TE=3.03 ms, 8 degree flip-angle, 1 slab, slice-matrix size=256×256, slice thickness=1 mm, field of view=256 mm, isotropic voxel-size=1.0×1.0×1.0 mm. Structural images were defaced by Schoffelen and colleagues. Preprocessing of the structural MRI was performed with Freesurfer (66), using the recon-all pipeline and a manual inspection of the cortical segmentations. Region-of-interest analyses were selected from the PALS Brodmann’ area atlas (67) and the Destrieux Atlas segmentation (68).

Functional images were acquired with a T2^*\**^-weighted functional echo planar blood oxygenation level dependent (EPI-BOLD) sequence. The full acquisition details, available in (36), are here summarized for simplicity: TR=2.0 seconds, TE=35ms, flip angle=90 degrees, anisotropic voxel size=3.5×3.5×3.0 mm extracted from 29 oblique slices. fMRI was preprocessed with fMRIPrep with default parameters (69). The resulting BOLD times series were detrended and de-confounded from 18 variables (the 6 estimated head-motion parameters (trans_*x,y,z*_, rot_*x,y,z*_) as well as the first 6 noise components calculated using anatomical CompCorr (70) and 6 DCT-basis regressors using nilearn’s clean_img pipeline and otherwise default parameters (71). The resulting volumetric data lying along a 3mm “line” orthogonal to the mid-thickness surface were linearly projected to the corresponding vertices. The resulting surface projections were spatially decimated by 10, and are hereafter referred to as voxels, for simplicity. Finally, each group of 5 sentences were separately and linearly detrended. Note that our cross-validation never splits such groups of five consecutive sentences between the train and test sets. Two subjects were excluded from the fMRI analyses because of difficulties processing the metadata, resulting in 100 fMRI subjects.

#### 4.2.3 Magneto-encephalography (MEG)

The MEG time series were pre-processed using MNE-Python and its default parameters except when specified (72). Signals were band-passed filtered between 0.1 and 40 Hz filtered, spatially corrected with a Maxwell Filter, clipped between the 0.01^*st*^ and 99.99^*th*^ percentiles, segmented between -500 ms to +2,000 ms relative to word onset and baseline-corrected before t=0. Reference channels and non-MEG channels were excluded from subsequent analyses, hence leading to 273 MEG channels per subject. We manually co-referenced (i) the skull segmentation of subjects’ anatomical MRI with (ii) the head markers digitized prior to MEG acquisition. A single-layer forward model was made. Because of lack of empty-room recordings, the noise covariance matrix used for the inverse operator was estimated from the zero-centered 200ms of baseline MEG activity preceding word onset. Subjects’ source space inverse operators were computed using a dSPRM. The average brain responses displayed in Figure 1D were computed as the square of the average evoked related field across all words for each subject separately, averaged across subjects and finally divided by their respective maxima, to highlight temporal differences. Video 1 displays the average sources without normalization. Seven subjects were excluded from the MEG analyses because of difficulties processing the metadata, resulting in 92 MEG subjects.

### 4.3 Noise Ceiling: Brain *→* Brain mapping

To estimate the amount of explainable signals in each MEG and each fMRI recordings, we trained and evaluated, through cross-validation, a linear mapping model *W* to predict the brain responses of a given subject to a each sentence *Y* from the aggregated brain responses of all other subjects who read the same sentence *X*. Specifically, five cross-validation splits were implemented across 5-sentence blocks with scikit-learn ‘GroupKFold’ (73). For each word of each sentence *i*, all but one subject who read the corresponding sentence were averaged with one another to form a template brain response: *x*_*i*_ *∈* ℝ^*n*^ with *n* the number of MEG channels or fMRI voxels, as well as a target brain response *y*_*i*_ *∈* ℝ^*n*^ corresponding to the remaining subject. *X* and *Y* were normalized (mean=0, std=1) across sentences for each spatio-temporal dimension, using a robust scaler clipping below and above the 0.01^*st*^ and 99.99^*th*^ percentiles, respectively. A linear mapping *W ∈* ℝ^*n×n*^ was then fit with a ridge regression to best predict *Y* from *X* on the train set:

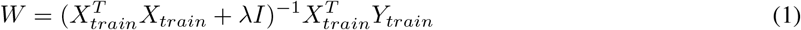

with *λ* the *l*2 regularization parameter, chosen amongst 20 values log-spaced between 10^*−*3^ and 10^8^ with nested leave-one-out cross-validation for each dimension separately. Brain predictions *Ŷ* = *WX* were evaluated with a Pearson correlation on the test set:

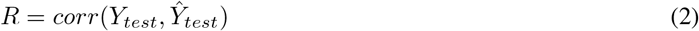

For the MEG source noise estimate, the correlation was also performed after source projection:

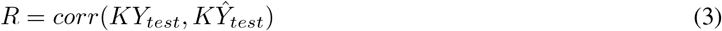

with *K ∈* ℝ^*n×m*^ the inverse operator projecting the *n* MEG sensors onto *m* sources. Correlation scores were finally averaged across cross-validation splits for each subject.

### 4.4 Similarity: Network *→* Brain mapping

To estimate the functional similarity between artificial neural networks and each brain, we followed the same analytical pipeline used for noise ceiling, but replace *X* with the activations of the deep learning models. Specifically, using the same cross-validation, and for each subject separately, we trained a linear mapping *W ∈* ℝ^*o,n*^ with *o* the number of activations, to predict brain responses *Y* from the network activations *X. X* was normalized across words (mean=0, std=1).

To account for the hemodynamic delay between word onset and the BOLD response recorded in fMRI, we used a finite impulse response (FIR) model with five delays (from 2 to 10 seconds) to build *X*^*\**^ from *X. W* was found using the same ridge regression described above, and evaluated with the same correlation scoring procedure. The resulting brain correlation scores measure the linear relationship between the brain signals of one subject (measured either by MEG or fMRI) and the activations of one artificial neural network (e.g a word embedding). For MEG, we simply fit and evaluate the model activations *X* at each time sample independently.

In principle, one may orthogonalize low-level representations (e.g. visual features) from high-level network models (e.g. language model), to separate the specific contribution of each type of model. This is because middle layers have access to the word-embedding layer, and can, in principle, simply copy some of its activations. Similarly, word embedding can implicitly contain visual information: e.g. frequent words tend to be visually smaller than rare ones. In our case, however, the middle layers of transformers were much better than word embeddings, and word embedding were much better than visual embeddings. To quantify the gain Δ*R* achieved by a higher-level model *M*_1_ (e.g. the middle layers of a transformer) and a lower level model *M*_2_ (e.g. a word embedding) we thus simply compare the difference of their encoding scores:

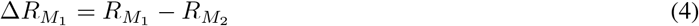

#### 4.4.1 Convergence analysis

All neural networks but the visual CNN were trained from scratch on the same corpus (cf. 4.1.2). We systematically computed the brain scores of their activations on each subject, sensor (and time sample in the case of MEG) independently. For computational reasons, we restricted model comparison on MEG encoding scores to ten time samples regularly distributed between [0, 2]s. Brain scores were then averaged across spatial dimensions (i.e. MEG channels or fMRI surface voxels), time samples and subjects to obtain the results in Figure 3. To evaluate the convergence of a model, we computed, for each subject separately, the correlation between (1) the average brain score of each network and (2) its performance or its training step. Positive and negative correlations indicate convergence and divergence respectively. Brain scores above 0 before training indicate a fortuitous relationship between the activations of the brain and those of the networks.

#### 4.4.2 Feature importance

To systematically quantify how the architecture, the accuracy and the learning of the artificial neural networks impacted their ability to linearly correlate with brain activity, we fitted, for each subject separately, a random forest across the models’ properties to predict their brain scores, using scikit-learn’s RandomForest (44; 73). Specifically, we input the following features to the random forest: the training task (causal language modeling vs. masked language modeling), the number of attention heads *∈* [4, 8], total number of layers *∈* [4, 8, 12], dimensionality *∈* [128, 256, 512], training step (number of gradient updates, *∈* [0, 4.5*M*], accuracy and the relative layer position of the representation (between 0 the first layer and 1 the last layer). The performance of the random forests was evaluated with a Pearson correlation *R* using a five-split cross-validation across models, for each subject separately.

“Feature importance” summarizes how each of the covarying properties of the models (their task, their architecture, etc) specifically impacts on brain scores. Feature importance is quantified with Δ*R*: the decrease in *R* when shuffling one feature (using 50 repetitions). For each subject, we reported the average decrease across the cross-validation splits (Figure 3). The resulting scores (Δ*R*) are expected to be centered around 0 if the corresponding feature does not impact brain score (even if it is indirectly correlated with it), and positive otherwise.

### 4.5 Population statistics

To estimate the robustness of our results, we systematically performed second-level analyses across subjects. Specifically, we applied Wilcoxon signed-rank tests across subjects’ estimates to evaluate whether the effect under consideration was systematically different from the chance level. The p-values of individual voxel/source/time samples were corrected for multiple comparison, using a False Discovery Rate (Benjamini/Hochberg) as implemented in MNE-Python. Error bars and *±* refer to the standard error of the mean (SEM) interval across subjects.

### 4.6 Ethics

This study was conducted in compliance with the Helsinki Declaration. No experiments on living beings were performed for this study. These data were provided (in part) by the Donders Institute for Brain, Cognition and Behaviour after having been approved by the local ethics committee (CMO – the local “Committee on Research Involving Human Subjects” in the Arnhem-Nijmegen region).

## 5 Acknowledgement

This work was supported by ANR-17-EURE-0017, the Fyssen Foundation, and the Bettencourt Foundation to JRK for his work at PSL.

## Footnotes

1 Algorithms were trained each on 8 GPUs using early stopping with training perplexity criteria, 16 streams per batch, 128 words per stream, epoch size of 200 000 streams, 0.1 dropout, 0.1 attention dropout, gelu activation, inverse (sqrt) adam optimizer with learning rate 0.0001, 0.01 weight decay.

2 https://github.com/attardi/wikiextractor

